# Modeling Electrocardiogram (ECG) Waveforms Using Group Theory and Symmetry Elements

**DOI:** 10.1101/2024.10.18.619135

**Authors:** Samuel J. Frueh

**Affiliations:** University of Connecticut, Institute of Materials Science

## Abstract

We present a what we believe to be a novel approach for generating electrocardiogram (ECG) waveforms using group theory and the algebraic structure of abstract symmetry elements from point groups up to order 10. Unlike traditional ECG modeling, which relies on physiological simulation or differential equations with predefined boundary conditions, our approach uses algebraic structures without predefined boundary conditions. This method allows us to explore the underlying algebraic symmetry of ECG signals, offering a perspective on biological signal representation. The flexibility and abstract nature of this framework opens potential applications for interdisciplinary research in mathematical biology, signal processing, and computational biology.

## 1. Introduction

The electrocardiogram (ECG) is a fundamental diagnostic tool for assessing cardiac function. Traditional approaches to modeling ECG signals involve either physiological models based on cardiac electrophysiology or purely mathematical approaches such as Fourier transforms, wavelets, or polynomial approximations. These methods, while effective, depend heavily on specific boundary conditionsand physical para meters that are intrinsic to the heart’s anatomy and physiology.

In recent years, however, the concept of **biological relativity** has gained attention as a framework for understanding complex biological systems. Proposed by Denis Noble, biological relativity posits that no single level of biological organization is privileged over others in determiningfunction and behavior. Rather, causation isdistributed across multiple levels, including molecular, cellular, tissue, and systemic levels, all interacting to contribute to the emergent properties of living organisms. This principle challenges reductionist views that prioritize one level of causation over others and instead emphasizes a more holistic approach.

Our work relates directlytotheidea of biologicalrelativity by employingan abstract mathematical representation,group the oretical algebra, to generate ECG-like waveforms. We illustrate that the emergent properties of the heart, as reflected in the ECG, can be described without privileging any specific biological level. Instead, these properties may emerge from algebraic relationships that embody the interplay between various underlying components. This perspective aligns with biological relativity, as it suggests that emergent behaviors, like the cardiac cycle, are not driven by one level of any one biological organization but rather are a product of interactions across all levels.

Group theory,widely used in chemistry and physics for understand molecular symmetry and particle dynamics, offers a powerful toolkit for exploring abstract relationships in dynamic systems. By applymg froup algebra and symmetry elements to mode_l_ ECG signala, we p owde an abstract, boundary-condition-free representation that captures the essential features of ECG waveforms, including the P-wave, QRS complex, and T-wave.

## 2. Methods

### 2.1 Group Theory and Symmetry Elements

We considered point groups of order up to 10, each represented by abstract elements (E, dA dB, dC… etc.). These elements follow algebraic group rules and are used to construct functions of time (E(t), dA(t), dB(t)… etc.) representing incremental changes in waveform components. The goal was to derive combinations of symmetry elements that resemble the ECG waveform in its key features.

### 2.2 Construction of Multiplication Tables

For each point group, multiplication tables are constructed to define the interaction of group elements. The multiplication results are represented in terms of functions involvingtime-dependent elements, allowing for the formation of waveform components that do not depend on physical boundary conditions or physiological constrai nts. This algebraic framework allows each component (P-wave, QRS com plex, T-wave) to be expressed through com binations of group elementsand corresponding funotions.

### 2.3 Function Assignments for ECG Components

- **P-wave:** The functions used to represent the P-wave take the form of Gaussian and sine-Gaussian functions, which generate a small rounded peak characteristic of the P-wave.
- **Q-wave:** This function is a negative Gaussian, capturing the small downward deflection of the Q-wave.
- **R-wave:** Modeled using a sharp Gaussian function that produces the prominent peak characteristic of the R-wave.
- **s-wave:** This function captures the negative deflection that follows the R-wave.
- **T-wave:** Modeled using a combination of functions, which are derived from point group. These functions represent a smooth, rounded peak that follows the QRS complex.

Ideal P,T, Q, R, S curves were numerically considered to be representationsforevery element of every symmetry group upto order 10. Group algebraic structure was used to determine the numerical structure of the curves that MUST follow for the symmetry elementE, and a function was fittedtothe curves for E(t). Subsequently, functions were determined for each element (dX(t))that must follow given the function for E(t) in every symmetry group up to order 10. This database of functions (of dX(t)) was searched to find the best fit forthe P T Q R S curves. Thus each curve is associated with the element or sum of elements of one or more point groups where E is represented by a function E(t) as shown above.

## 3. Results

The generated ECG waveform is a combination of the P-wave, QRS complex, and T-wave, constructed using abstract group elements without any imposed boundary conditions. The resulting wave form captures the essential features of a standard ECG, with flexibility in adjusting the functions to match the desired characteristics of each component. The abstract nature of the construction meansthat the wave form is not tied to specific physiological constraints, allowing for a generalized representation.

Residual analysis between the model waveform and a real ECG waveform showed promising fidelity, particularly in capturing the sharp features of the QRS complexand the overall timing of the signal. The ability to generate these waveforms without boundary conditions suggests that the underlying structure of biological waveforms could be interpreted through algebraic relationships and symmetry.

P-wave

- Point Group: *C*_2_, *D*_1_
- Symmetry Elements: *dA, dB*
- Function *E(t): E*_3_*(t),E*_5_*(t)*

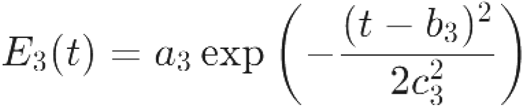

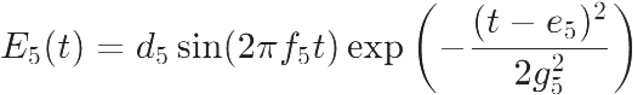

Here are the specific forms of the functions for the Q-wave:
- **Point Group:** *C*_4_
- **Symmetry Elements:** *dC*
- **Function *E***_7_***(t)*** This function is used to represent the Q-wave, which is typically a small negative deflection before the R-wave. The form could be:

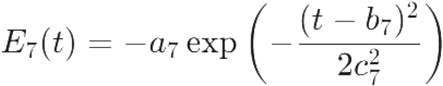

where *a*_7_ *b*_7_, and *c*_7_ are parameters that control the amplitude, center, and width of the Gaussian function. This negative exponential shape captures the brief downward dip characteristic of the Q-wave in an ECG.

Here are the specific forms of the functions for the R-wave:

- **Point Group:** *C*_6_
- **Symmetry Elements:** *dA*
- **Function** *E*_10_(t): This function represents the R-wave, which is the most prominent peak in the ECG waveform. The form can be modeled using a sharp Gaussian function:

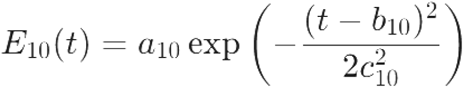

where *a*_10_, *b*_10_, and *c*_10_ are parameters that control the amplitude, the center (representing the timing of the R-wave peak), and the width of the peak (with a small value for *c*_10_ to create a sharp peak characteristic of the R-wave).

Here are the specific forms of the functions for the T-wave:

- **Point Group:** *C*_3_
- **Symmetry Elements: *dA, dC***
- **Functions** *E*_18_, (*t*): *E* _20_ *(t)* : The T-wave represents **a** rounded peak following the QRS complex. It can be modeled by combining two functions to produce a smooth shape:

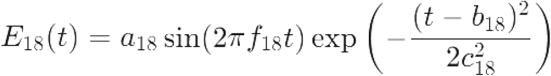
- Where *a*_18_, *f*_18_,*b*_18_, and *c*_18_ control the amplitude, frequency, center, and width of the sine-Gaussian function.

Additionally, *E*_20_(*t*) may take the form:

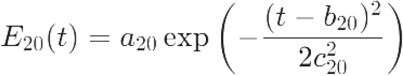

where *a*_20_, *b*_20_, and *c*_20_ control the amplitude, center, and width. This combination of functions captures the broad and smooth characteristics of the T-wave.

Here are the specific forms of the functions for the S-wave:

- **Point Group:** *D*_3_
- **Symmetry Elements: *dB***
- **Function** *E*_12_*(t):*This function represents the S-wave, which is a negative deflection following the R-wave. The form could be:

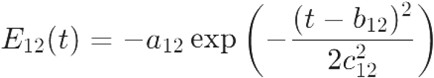

where *a*_12_, *b*_12_and *c*_12_ are parameters that determine the amplitude, center, and width of the S-wave. The negative sign ensures the downward characteristic of the S-wave is properly represented.

↓

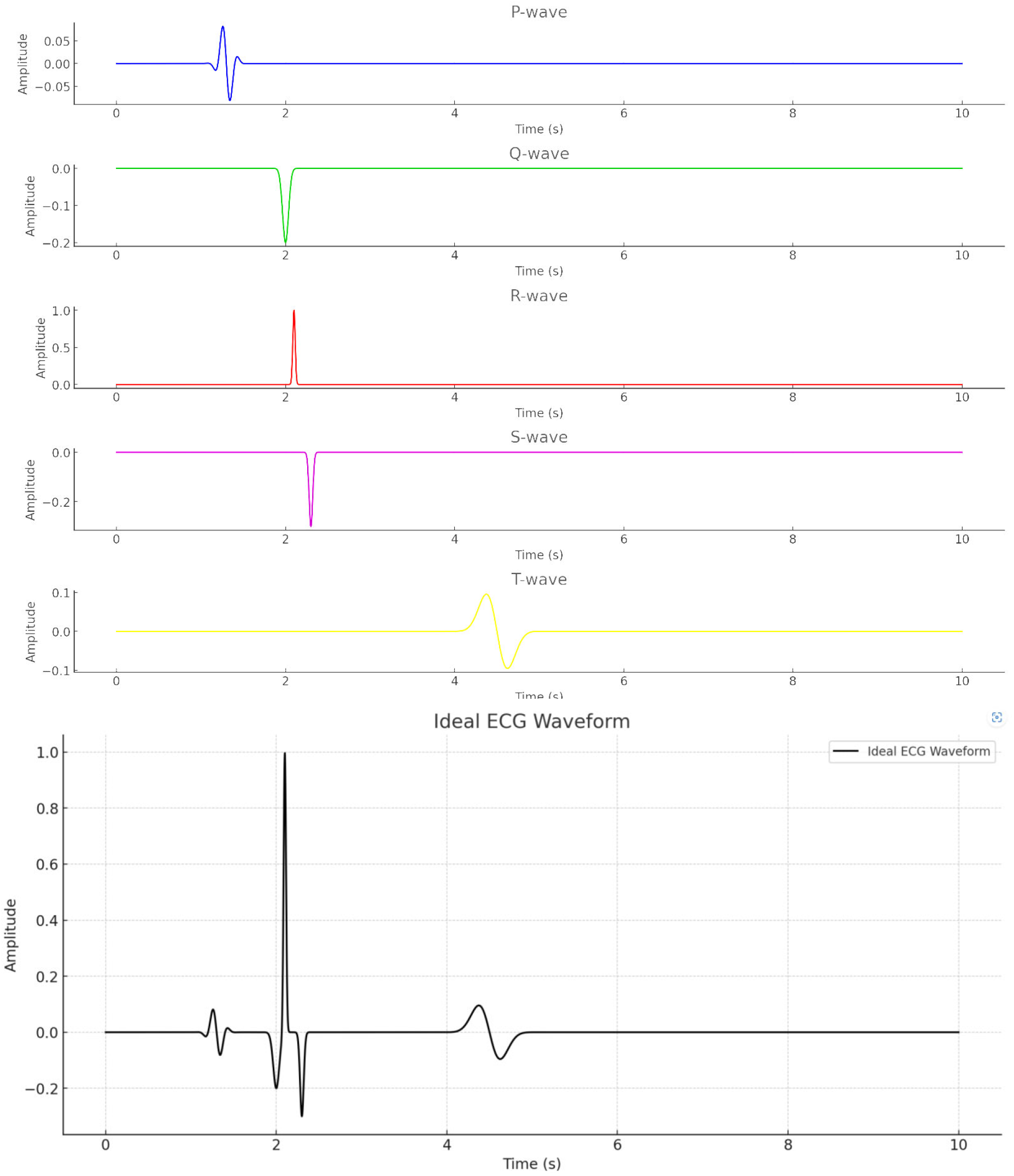

## 4. Discussion

This work represents a new direction in ECG modeling by using group theory to derive waveform features without the constraints of physiological modeling. The algebraic structure provides a flexible framework for creating a wide range of waveforms, suggesting potential applications in signal processing, synthetic data generation, and the exploration of general biological rhythms. By abstracting the ECG to group algebra, we open new opportunities for interdisciplinary collaboration, linking mathematical biology, abstract algebra, and computational modeling.

### 4.1 Non-Privileged Levels of Causation, Biological Relativity, and Gödel’s Incompleteness Theorem

The concept of non-privileged levels of causation implies that no single level of description (such as cellular, molecular, or systemic) has absolute precedence over another in determining the behavior of a biological system. This idea suggests that emergent properties of biological systems are influenced by multiple interdependent levels rather than being driven solely by any specific component in this context, our approach to modeling the ECG using group theory challenges the not on that the physiological characteristics of the heart are the only valid perspective for understanding the dynamics of cardiac signals.

By employing an abstract algebraic structure, we suggest that the emergent features of the ECG may be derived from purely mathematical relationships, highlighting the interplay between different levels of causation. This approach provides an alternative to the reductionist view, where complex biological behaviorsare often reduced to the function of individual components. Instead, our model emphasizes a holistic perspective where the underlying symmetry relationships play a significant role in shaping the observed ECG waveform. This aligns with the concept of biological relativity, which asserts that causation is distributed across all levels of biological organization, and that no single level holds a privileged position.

It is important to note, however, that the ECG waveform generated here was matched to a known waveform, rather than being de rived a priori. This limitation indicates that while our model does not require traditional boundary conditions, it still relies on matching to empirical data. Critics might argue that the functions and even the point groups themselves could be considered boundary conditions in disguise, as they provide structure and constraints to the generated waveform. In response to this criticism, we propose that a true test of the model’s validity would involve generating the ECG waveform entirely from first principles, without reliance on empirical matching. We also note that the failure to derive the ECG waveform from first principals, and/or given a proven inability to do so, aligns with the context that sufficiently complex systems cannot be fully described by a finite set of axioms.

To settle this argument, future work could focus on deriving the form of from fundamental physiological principles, rather than empirical fitting. Additionally, it would be essential to explore whether the unused symmetry elements, once characterized, can predict physiological phenomena that are currently unmeasured or poorly understood. This would provide stronger evidence that the algebraic framework captures inherent aspects of cardiac activity beyond the conventional ECG.

Furthermore, the concept of non-privileged levels of causation resonates with the notion that multiple descriptions—be they physiological, mathematical, or even quantum—could provide complementary insights into the heart’s function. The use of group theory thus opens up the possibility of viewing cardiac activity not just through the lens of physiology, butalso through the lens of abstract algebra, potentially revealing hidden symmetries that contribute to the emergent properties of the heartbeat.

By employing an abstract algebraic structure, we suggest that the emergent features of the ECG may be derived from purely mathematical relationships, highlighting the interplay between different levels of causation. This approach provides an alternative to the reductionist view, where complex biological behaviors are often reduced to the function of individual components. Instead, our model emphasizes a holistic perspective where the underlying symmetry relationships play a significant role in shaping the observed ECG waveform.

### 4.2 Conjecture on Unused Symmetry Elements and Physiological Phenomena

An intriguing aspect of this model is the presence of unused symmetry elements and functions that were not required for constructing the primary features of the ECG waveform. We conjecture that these unused elements may correspond to real physiological phenomena that contribute to the heartbeat but are not directly captured in an ECG signal. These phenomena could include:

- **Mechanical Movements of the Heart Wall:** While the ECG captures the electrical activity of the heart, mechanical deformations and movements of the heart wall are not directly represented. The unused symmetry elements might correspond to the forces and stresses experienced duringthe contraction and relaxation of the heart muscle, which contribute to efficient pumping but are not part of the electrical signal.
- **Autonomic Nervous System Influences:** The influence of the autonomic nervous system (ANS) on heart rate variability and myocardial contractility is a complex interplay that affects the overall cardiac cycle. The unused symmetry elements might represent aspects of sympathetic and parasympathetic regulation that do not generate direct electrical signals but modulate the amplitude and timing of those signals.
- **Subthreshold Electrical Activities:** Not all electrical activities reach the threshold needed to produce visible deflections in an ECG. These subthreshold events could be linked to the unused symmetry operations, contributing to the background electrical environment of cardiac cells that ensures the heart’s readiness for activation.
- **Vascular Coupling:** The interaction between the cardiac muscle and the surrounding vasculature (e.g., coronary perfusion) affects the efficiency of the heartbeat. The unused symmetry elements might correspond to vascular coupling effects that are not evident in surface ECG recordings but still influence cardiac function.

However, it is also important to consider a critical perspective on this conjecture. One possible criticism is that the assumption that every unused symmetry element must correspond to a physiological phenomenon could be unfounded. It may be that these unused elements are artifacts of the mathematical framework rather than reflections of real physiological processes. If this criticism were valid, it would imply that the group-theoretic approach might not have a one-to-one correspondence with biological reality. I nstead.it could suggest that the mathematical model, while useful, may over fit or misrepresent aspects of the system.

If these unused elements do not have physiological correlates, it could mean that the group theory model captures a broader, more abstract set of relationships that extend beyond the physiological specifics of the heart. This would imply that the model is not entirely bound to the real-world mechanics of cardiac function and may instead represent potential or hypothetical dynamics that do not manifest in observable physiology.

To address this criticism and settle the argument, further experimental validation is necessary. Specifically,we would need to determine whether the unused symmetry elements predict measurable physiological effects. For instance:

- **Targeted Experiments:** Conduct targeted experiments to measure potential physiological features that are hypothesized to correspond to the unused symmetry elements. This could include imaging mechanical heart movements with high resolution or using advanced electrophysiological techniques to detect subtle subthreshold activities.
- **Correlation Studies:** Explore correlations between unused symmetry elements and physiological metricsthat are nottypically captured in standard ECG recordings. This could involve simultaneous measurements of heart rate variability, cardiac wall motion, and vascular dynamics, compared against predictions made by the group-theoretic model.
- **Model Refinement:** If no physiological correlates are found, it may be necessary to refine the model, either by constraing the group structure or by modifying the function assignments to better align with observed biological behavior. This could involve introducing boundary conditions that selectively constrain which elements are used in constructing the ECG waveform.

Ultimately, determining whether unused symmetry elements correspond to real physiological phenomena will require both theoretical exploration and experimental verification. If proven valid, this would significantly enhance our understanding of the hidden complexities of cardiac function and suggest that biological rhythms may be governed by deeper, previously unrecognized symmetries. Conversely, if the criticism holds, it will serve as an opportunity to refine the model to ensure that it more accurately reflects the physiological realities of the heart.

### 4.3 Experimental Testing and Measurement

To test the validity of this conjecture, new experimental approaches could be devised to measure physiological features that might correspond to the unused symmetry elements:

- **Cardiac MRI and Echocardiography:** Imaging techniques like cardiac MRI or echocardiography could be used to measure the mechanical deformation of the heart wall, potentially correlating unused symmetry elements with specific mechanical features of cardiac motion.
- **Heart Rate Variability (HRV) Analysis:** The influence of the autonomic nervous system on cardiac activity could be examined using HRV analysis. By comparing HRV metrics with the unused symmetry elements, it may be possible to identify links between group-theoretic elements and autonomic modulation.
- . **Intracardiac Electrophysiology Studies:** Subthreshold electrical activities could be investigated through invasive electrophysiology studies, where intracardiac electrodes are used to detect signals that do not appear in surface ECGs. These subthreshold signals might correspond to the unused algebraic elements in our model.
- **Coronary Flow Measurements:** The relationship between vascular dynamics and cardiac electrical activity could be examined using coronary flow measurements during different phases of the cardiac cycle, identifying potential connections with unused symmetry elements.

## 5. Conclusion

We have demonstrated the feasibility of generating an ECG-like waveform using group theory and abstract algebraic elements. This approach diverges from traditional physiological models by focusingon symmetry and algebraic structure, offering a unique perspective on biological signal generation. By considering non-privileged levels of causation and Godel’s incompleteness theorem, we suggest that the unused symmetry elements in our model may represent real but currently unmeasured physiological phenomena. Future work will focus on introducing boundary conditionsto make the model more applicable to real-world scenarios, exploring the physiological correlates of unused symmetry elements, and applying this framework to other physiological signals.

## Acknowledgments

We acknowledge the contributions of interdisciplinary research that inspired the use of group theory in biological signal modelingand the potential implications for future studies in mathematical biology.

